# Towards Interpretable Multitask Learning for Splice Site and Translation Initiation Site Prediction

**DOI:** 10.1101/2023.10.16.562631

**Authors:** Espoir Kabanga, Arnout Van Messem, Wesley De Neve

## Abstract

In this study, we investigate the effectiveness of multi-task learning (MTL) for handling three bioinformatics tasks: donor splice site prediction, acceptor splice site prediction, and translation initiation site prediction. As the foundation for our MTL approach, we use the SpliceRover model, which has previously been successful in predicting splice sites. While providing benefits such as efficient resource utilization, reduced complexity, and streamlined model management, our findings show that the newly introduced MTL model performs comparably to the SpliceRover model trained separately for each task (single-task models), with a slight decrease in specificity, sensitivity, F1-score, and Matthews Correlation Coefficient (MCC). However, these differences are statistically insignificant (the specificity decreased with 0.0081 for acceptor splice site prediction and the MCC decreased with 0.0264 for TIS prediction), emphasizing the comparable performance of the MTL model. We further analyze the effectiveness of our MTL model using visualization techniques. The outcomes indicate that our MTL model effectively learns the relevant features associated with each task when compared to the single-task models (presence of nucleotides with a higher contribution to donor splice site prediction, polypyrimidine tracts in the upstream of acceptor splice sites, and the Kozak sequence). In conclusion, our results show that the MTL model generalizes well across all three tasks.

## 1 Introduction

Over the last decade, machine learning research in the field of computational biology has gained in popularity, primarily driven by the application of deep learning techniques to several biological tasks [1]. The effectiveness of these techniques heavily relies on data quantity and quality, which researchers consider to be two major challenges, mainly due to privacy, security, and cost constraints [2]. One way to address these challenges is to build deep learning models that can learn simultaneously from different but closely related biological tasks. These models are known as multi-task learning (MTL) models [3].

MTL simultaneously trains a model to learn multiple tasks, utilizing a shared representation to optimize learning across all tasks [4]. This approach has achieved significant success in computer vision, medical imaging, speech recognition, and natural language processing [5, 6, 7, 8]. In MTL, hard parameter sharing and soft parameter sharing are two different ways to share information between tasks. Hard parameter sharing involves using a single model with shared parameters for all tasks, while soft parameter sharing uses separate models for each task with regularization to encourage parameter similarity. Hard sharing assumes all tasks are similar, while soft sharing allows for task-specific and shared parameters [9]. MTL is of particular interest in computational biology due to the noisy and limited availability of genomic data, enabling faster and more generalizable learning across multiple tasks [3, 10].

The prediction of splice sites plays a critical role in gene expression [11]. Deep learning techniques have considerably advanced splice site prediction, showing substantial improvements in false discovery rate [12] and accuracy for human genomic data [13]. Additionally, they have demonstrated the ability to identify conserved genomic elements and annotate splice sites in new genomes by selecting suitable models based on taxonomic similarity [14].

Correct identification of translation initiation sites (TIS) is crucial for discovering gene structure and regulating gene expression. TIS detection relies on the Scanning model [15], which identifies the first AUG codon at the 5’ end of an mRNA transcript in eukaryotes as a valid TIS . Machine learning techniques have been employed for automated TIS annotation, with recent deep learning approaches, such as TITER [16] and TISRover [17], demonstrating high effectiveness. Accurate TIS prediction is essential in computational biology for understanding gene expression and regulation.

## 2 Data and methods

### 2.1 Dataset description

Our experiments involved three distinct prediction tasks: (1) donor splice site prediction, (2) acceptor splice site prediction, and (3) TIS prediction. For this purpose, we used a dataset consisting of DNA sequences obtained from the *Arabidopsis thaliana* genome. Each task was associated with a balanced dataset that included an equal number of confirmed true (positive) and confirmed false (negative) candidate sites. Specifically, for each task, the dataset comprised 11,520 DNA sequences for training, 1,280 DNA sequences for testing, and 3,200 sequences for validation. Within each training, validation, and test partition, half of the sequences contained positive candidate sites and the remaining half contained negative candidate sites. For donor splice site prediction and acceptor splice site prediction, we used a portion of the dataset that was employed in [18], with each sequence consisting of 602 nucleotides. For TIS prediction, we utilized a portion of the dataset employed in [19], with each sequence consisting of 300 nucleotides.

### 2.2 Data preprocessing

To ensure consistency across all three prediction tasks, we made modifications to the dataset used for donor splice site and acceptor splice site prediction. Initially, these datasets consisted of sequences with a length of 602 nucleotides, with the candidate site (a GT pair for donor splice sites and an AG pair for acceptor splice sites) located in the middle of the sequence, specifically at positions 301 and 302. In contrast, the dataset for TIS prediction had sequences with a length of 300 nucleotides, with the candidate site (an ATG triplet) located in the middle of the sequence, at positions 150-152.

In order to standardize the length of the sequences across all three prediction tasks, we trimmed 151 nucleotides from the beginning and end of the sequences used for donor splice site and acceptor splice site prediction. This resulted in sequences with a length of 300 nucleotides, the same as those used for TIS prediction.

To prepare the dataset for our different models, the DNA sequences were converted into a one-hot encoding format. In this encoding scheme, each nucleotide is represented as a vector of four binary values, with A being represented as [1,0,0,0], C as [0,1,0,0], G as [0,0,1,0], and T as [0,0,0,1]. This conversion is a crucial step in the preprocessing of the data as it enables the network models to better understand the sequence information and identify relevant features in the input data.

### 2.3 Model description

To construct our MTL model, we made use of the SpliceRover model, as described in [12]. We kept all the parameters identical, such as the number of convolutional layers, the number of filters and their sizes, and the number of neurons in the fully-connected layer. The only change we made was in the optimizer, where we replaced stochastic gradient descent with the Adam optimizer [20], which has the advantage of using adaptive learning rates. We then trained the resulting model on all three tasks simultaneously, following a soft parameter learning approach.

Figure 1 provides a visual representation of our MTL model architecture. We adopted one input layer for each task, thus coming with three input layers in total. Moreover, for each task, we included a task-specific layer for feature learning, consisting of four convolutional layers. The first two convolutional layers were each followed by a dropout layer (p=0.2), while the other two convolutional layers were each followed by a max pooling layer with pool sizes three and four, respectively, and a dropout layer (p=0.2). The parameters of the four task-specific layers were the same as those in the SpliceRover model. Following the task-specific layers, we added a concatenate layer to combine the features learned separately from each task. The concatenation operation allows the model to increase the feature space and facilitate feature interaction between tasks. The combined features were then passed through shared convolutional layers, followed by a max pooling layer (pool size=4) and a dropout layer (p=0.2). The parameters of the shared convolutional layer were the same as those of the fifth convolutional layer in the SpliceRover model. The shared layers reduce the risk of overfitting, and the model is more likely to find a representation that captures all tasks [3].

**Figure 1:**
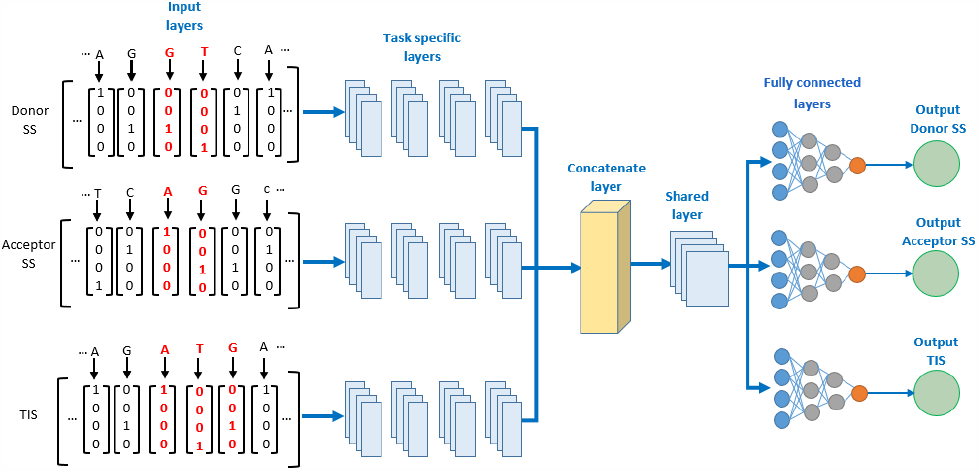
Overview of model architecture.

The model had three output layers, one for each task, and each output layer consisted of two neurons, one for generating a positive prediction and another one for generating a negative prediction.

### 2.4 Evaluation metrics

To assess the effectiveness of our MTL model, we used four distinct evaluation metrics: specificity, sensitivity, F1-score, and Matthews Correlation Coefficient (MCC).

Specificity assesses accuracy in identifying negative cases, while sensitivity measures the accuracy for positive cases. F1-score is a balanced measure of precision (accuracy of positive predictions) and recall (completeness of positive predictions). The MCC takes into account true and false positives and negatives for a thorough evaluation of binary classification.

## 3 Results

To account for the random initialization of weights, we conducted five training runs and calculated the mean values for the different performance metrics used. Each training session consisted of a maximum of 50 epochs. To prevent overfitting, we used an early-stopping approach with a patience of five. This means that training stops if the validation loss of a model does not decrease after five epochs. Since we were addressing three tasks, we had three distinct losses, one for each task. The overall loss of our MTL model was determined by computing the uniformly weighted sum of the individual task losses. Our findings indicate that the proposed MTL model performs similarly to the single-task models, across all the tasks and for each of the performance metrics (differences are statistically insignificant). Detailed results can be found in Table 1.

**Table 1:** Performance comparison of the mean and standard deviation of the single-task models and the MTL model.

|  |  | Specificity | Sensitivity | F1-score | MCC |
| --- | --- | --- | --- | --- | --- |
| Donor SS | single-task model | 0.9367±0.0121 | 0.9434±0.0073 | 0.9403±0.0028 | 0.8803±0.0064 |
| | MTL model | 0.9326±0.0064 | 0.9315±0.0047 | 0.9320±0.0030 | 0.8642±0.0062 |
| Acceptor SS | single-task model | 0.9265±0.0043 | 0.9361±0.0049 | 0.9316±0.0049 | 0.8627±0.0096 |
| | MTL model | 0.9184±0.0150 | 0.9269±0.0167 | 0.9229±0.0029 | 0.8456±0.0045 |
| TIS | single-task model | 0.9199±0.0189 | 0.9211±0.0136 | 0.9206±0.0033 | 0.8413±0.0076 |
| | MTL model | 0.9103±0.0117 | 0.9044±0.0162 | 0.9070±0.0038 | 0.8149±0.0052 |

In comparison with existing models in terms of F1-score, we observed similar trends in the performance of our MTL model. Specifically, our MTL model showed a slight improvement over SpliceFinder [13], with a margin of 0.0098 and 0.0043 for donor and acceptor splice sites, respectively. Moreover, when compared to EnsembleSplice [18], which outperforms other models presented in the literature and which was trained on the same dataset, the performance of the MTL model was slightly lower, with a difference of 0.0144 (donor splice sites) and 0.0232 (acceptor splice sites) in terms of F1-score. Regarding TIS prediction, our MTL model showed a decrease of 0.0186 in terms of F1-score, compared to TISRover [17] trained on our TIS dataset.

However, of higher interest is understanding whether our MTL model is able to concurrently learn the features that are specific to each task. To that end, we utilized the saliency maps approach presented in [21] to generate the contribution scores of each nucleotide in a DNA sequence. In particular, we computed the gradients of the output of a model with respect to its input. These gradients, also known as saliency values, can then be linked to each nucleotide in the DNA sequence under consideration, forming a saliency map [22]. To make the values more comparable across different sequences, we divided each saliency value by the average of the absolute values of all the saliency scores in the sequence. Furthermore, the average saliency was scaled down by a factor of 100 to provide a standard for normalization.

The sequence logos presented in Figure 2, generated using Logomaker [23], show the contribution (importance) score of each nucleotide at specific positions plotted along the *x*-axis. In the top row, both the single-task model and the MTL model for donor splice sites capture the common CAGGTAAGT pattern and exhibit similar contribution patterns. For acceptor splice sites in the middle row, both models learn the polypyrimidine tract, which is characterized by C and T nucleotides, upstream of the acceptor site. Regarding the TIS in the bottom row, both models identify similar patterns, with higher contribution scores for `G` and `C` at positions 3 and 4 downstream of TIS, and `A/G` (purine) at position -3 upstream of TIS, which aligns with the typical context described by the Kozak sequence [15]. While the sequence logos of both single-task and MTL models show similar contribution score magnitudes overall, the contribution score magnitudes of the single-task model for acceptor splice site prediction and the single-task model for TIS prediction are higher compared to the contribution score magnitudes of the MTL model.

**Figure 2:**
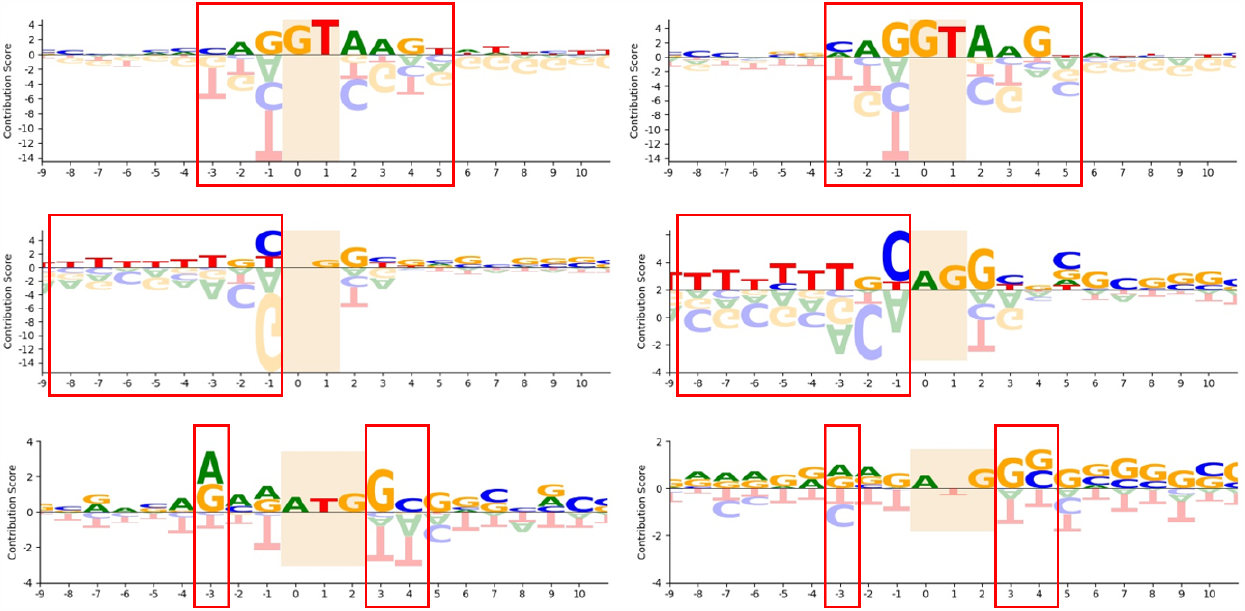
Comparison of sequence logos between the single-task models (left) and our MTL model (right) for predicting donor splice sites (top), acceptor splice sites (middle), and translation initiation sites (bottom).

## 4 Conclusions

We investigated the effectiveness of MTL for three bioinformatics tasks: donor splice site prediction, acceptor splice site prediction, and TIS prediction. Comparing our MTL model to the single-task models, we found that the proposed MTL model performed similarly without any statistically significant differences. In addition, our MTL model successfully learned the relevant patterns associated with all three tasks simultaneously, including the presence of nucleotides with a higher contribution to donor splice site prediction, polypyrimidine tracts in the upstream of acceptor splice sites, and the Kozak sequence for TIS. Furthermore, it is worth mentioning that our MTL model enables multiple predictive tasks to be handled within a single framework, resulting in lower computational costs, simplified deployment, and easier maintenance workflows, while still achieving comparable performance across the various predictive tasks.

In future work, we plan to investigate the impact of positive and negative transfer between tasks and compare MTL models with fine-tuned foundation models.

## Conflicts of interest

All authors declare that they have no conflicts of interest.

## Notes

### Competing Interest Statement

The authors have declared no competing interest.

